# Acquisition of Oral Microbiota is Driven by Environment, Not Host Genetics

**DOI:** 10.1101/2020.01.06.896126

**Authors:** Chiranjit Mukherjee, Christina O. Moyer, Heidi M. Steinkamp, Shahr B. Hashmi, Xiaohan Guo, Ai Ni, Eugene J. Leys, Ann L. Griffen

## Abstract

The human oral microbiota is acquired early in an organized pattern, but the factors driving this acquisition are not well understood. Microbial “heritability” could have far-reaching consequences for health, yet no studies have specifically examined the fidelity with which the oral microbiota are passed from parents to offspring. Some previous studies comparing monozygotic (MZ) and dizygotic (DZ) twins had suggested that host genetics has a role in shaping oral microbial communities, and also identified so called “heritable” taxa. However, these findings are likely to be confounded by shared environmental factors resulting from the well-established greater behavioral similarity among MZ twins. In addition, MZ and DZ twins share an equal portion of their parent’s genome, and so this model is not informative for studying direct parent to offspring transmission.

To specifically examine the contribution of genetics to the fidelity of transmission of bacteria from parents to offspring, we used a novel study design comparing fraction of shared species and strains between our genetically related group consisting of children and their biological mothers, with that of children and their adoptive mothers, constituting our genetically unrelated group. Fifty-five biological and 50 adoptive mother-child pairs were recruited along with 23 biological fathers and 16 siblings. Subjects were carefully selected to ensure the two groups were matched on child’s age. Three distinct habitats within the oral cavity: the saliva/soft tissue surface, supragingival biofilm, and subgingival biofilm, were sampled to comprehensively profile the oral microbiome. Our recently developed strategy for subspecies level characterization of bacterial communities by targeted sequencing of the ribosomal 16-23S intergenic spacer region (ISR) was utilized in the present study to track strain sharing between subjects, in addition to 16S rRNA gene sequencing for species analysis.

Results showed that oral bacterial community profiles of adoptive and biological mother-child pairs were equally similar, indicating no effect of host genetics on the fidelity of transmission. This was consistent at both species and strain level resolutions, and across all three habitats sampled. We also found that all children more closely resembled their own mother as compared to unrelated women, suggesting that contact and shared environment were the major factors shaping the oral microbiota. Individual analysis of the most abundant species also did not detect any effect of host genetics on strain sharing between mother and child. Mother-child strain similarity increased with the age of the child, ruling out early effects that are lost over time. No effect on the fidelity of mother-child strain sharing from vaginal birth or breast feeding was seen. Analysis of extended families showed that fathers and mothers were equally similar to their children. Cohabitating couples showed even greater strain similarity than mother-child pairs, further supporting the role of age, contact and shared environment as determinants of microbial similarity.

Based on these findings we suggest that the genetic effects on oral microbial acquisition observed in twin studies are more likely the result of confounding environmental factors based on greater behavioral similarity among MZ twins. Our findings suggest that these host mechanisms are universal to humans, since no effect of genetic relatedness on fidelity of microbial transmission could be detected. Instead, our findings point toward contact and shared environment being the driving factors of microbial transmission, with a unique combination of these factors ultimately shaping a highly personalized human oral microbiome.

## INTRODUCTION

The last decade has seen rapid growth in understanding the role of the human microbiome in health and disease. Foundational studies across multiple body sites have shown that the adult human microbiota consists of a shared, limited set of niche-specific species^1,2^. Our previous work exploring the assembly of oral microbiota from birth through the first year of life showed that the early childhood microbiota is acquired in an ordered sequence, with common oral species shared between children and mothers^3^. At the strain level, however, each individual harbors a highly personalized microbiota^4,5^. Role of intrinsic and extrinsic factors in acquisition of this individualized microbiome are yet not well understood. Given that strains have been found to be shared among family members^6^, a possible intrinsic factor influencing microbiota acquisition could be host genetics shared between parent and offspring.

A number of previous investigations into the interactions of microbes and host genetics have compared microbial communities of monozygotic (MZ) twins, and dizygotic (DZ) twins. Two of these studies have shown the oral microbiota of MZ twins to be slightly more similar than that of DZ twins, suggesting that host genetics influences microbial community composition^7,8^. These studies also identified taxa that are more likely to be “heritable”. Microbial “heritability” could have far-reaching consequences for health, since the inheritance of a dysbiotic community could confer increased risk for diseases with a microbial etiology. Yet no studies have specifically examined the fidelity with which the human oral microbiota are passed from parents to offspring.

A major deterrent to studying the role of shared genetics in parent to child microbial transmission has been the lack of a suitable case-control model. Both MZ and DZ twins share an equal portion of their parent’s genome and are therefore not informative for studying direct parent to offspring transmission. To specifically examine the contribution of genetics to the fidelity with which microbial species and strains are passed from parents to offspring, we used a novel study design comparing genetically related and unrelated mother-child pairs. The fraction of shared species and strains between children and their biological mothers, who share half of their genomes, was compared with that of children and their adoptive mothers, who had no genetic relationship. Extensive metadata was collected to examine potential confounders such as breastfeeding and birth mode. Given that humans share a core oral microbiota at the species level^9^, subspecies or strain level techniques are required to accurately track microbial transmission between individuals. We have developed a high-throughput strategy for sub-species/strain characterization of bacterial communities by targeting the ribosomal intergenic spacer region (ISR), and this approach has revealed highly personalized profiles among adults^5^. We employed this approach for sub-species analysis, along with 16S rRNA gene sequencing for species analysis. Microbial community composition is known to vary among distinct niches within the oral cavity that are important in oral disease. Therefore, to comprehensively profile the oral microbiota, we sampled three distinct oral habitats - soft tissue and saliva, and both supragingival and subgingival plaque biofilm.

Using this multi-habitat, multi-resolution approach, we comprehensively profiled oral microbial communities in parents and children. We detected no effect of genetic relationship on fidelity of transmission at either the species or strain level, and no effect in any of the three biologically important oral niches. Our findings suggest that contact and shared environment, not genetics, determine the transmission of oral microbes. An extended dataset showing high similarity between spouses further supported this observation.

## METHODS

### Inclusion/exclusion criteria, exam and sampling

Adoptive and biological mother-child dyads were enrolled to allow determination of the effect of genetic relatedness on the fidelity of oral bacterial transmission. IRB approval was obtained for this study, parents provided written consent and children over 7 years of age provided assent. Adoptive mother-child dyads were recruited through adoption agencies. The biological group was recruited to match the adoptive group on child’s age, and parent’s socioeconomic status. Only children adopted immediately at birth and unrelated to the adoptive family were included to minimize transmission of bacteria from the biological mother. Only genetic birth mothers were included in the biological group, and fathers and siblings from this group were also sampled when available. For both biological and adoptive groups, the minimum age for children was 3 months to allow the establishment of an oral bacterial community, while twelve years of age was the maximum. Exclusion criteria for all subjects were chronic disease affecting the oral cavity or immune system, and/or early onset periodontitis.

### Examination and history

A dental exam was conducted and recorded for both children and mothers that included caries, gingivitis, periodontitis, and plaque levels. A medical and social history was also obtained, including breastfeeding history and delivery mode.

### Sample Collection and Processing

Sampling was conducted at least 1 hour after home oral hygiene or consuming food or drink. Saliva and soft tissue were sampled with a Copan swab placed in the right lingual vestibule for 30 seconds and then swabbed into both buccal vestibules and across the tongue. Sterile microbrushes were used to collect supragingival plaque from the buccal surfaces of all teeth in the mandibular right quadrant, and sterile paper points were then inserted into the mesial sulcus of each of these teeth to obtain subgingival samples. Samples were placed separately in labelled tubes containing 200 µL ATL buffer (Qiagen, USA) and stored under refrigeration until transported to the lab for storage at -20°C. Bacterial genomic DNA was extracted using QIAamp DNA Mini Kit (Qiagen, USA), using an optimized protocol as described in Mukherjee et all 2018^5^.

### Sequencing Library Preparation

Two sets of sequencing libraries were prepared, targeting the 16S V1-V3 and 16-23S ISR, as described in Mukherjee et al 2018. Briefly, both the protocols are based on an optimized Illumina 16S Metagenomic Sequencing Library Preparation protocol (Illumina, USA), a two-step process where the target region is PCR amplified with gene-specific primers and then indexing barcodes are added through a second round of amplification. An optimization was made to the molecular protocol for the 16-23S ISR library such that only the ISR fragment was amplified as target, excluding the adjacent 16S region. This was achieved using the rD1f primer (5’-GGCTGGATCACCTCCTT)^10^ in place of the 1237F primer in the original protocol. The 16S V1-V3 libraries were sequenced on the Illumina MiSeq Platform with 300 base pair paired-end chemistry. The ISR libraries were sequenced on the Illumina HiSeq 2500 platform, with 250 base pair paired-end chemistry.

### Sequence data processing & taxonomic assignment

Demultiplexed reads from both the 16S V1-V3 library were processed as described in library Mukherjee et al. 2018^5^. Briefly, paired reads from the 16S library were merged using Mothur^11^, quality filtered using Mothur and custom Python scripts, and then mapped against our oral bacteria16S reference database CORE^12^ to generate species-level OTU abundance tables.

For the ISR library, denoising and sample inference was performed using DAD2^13^ version 1.10.0 for the unpaired read1s, as described in Mukherjee et al. 2018^5^, with quality filtering parameters optimized to suite this library. Specifically, parameters used in DADA2 processing protocol were adjusted for the primers used in this study and overall library quality (trimLeft=17, maxN=0, truncQ=2). Chimeric reads were removed with DADA2 using the function *removeBimeraDenovo* (method=“consensus”) and low throughput (<5000 input sequences) samples were excluded from the analysis. ISR ASVs present in only 1 sample were removed as suspect PCR artefacts. The detailed denoising pipeline in R is available at https://github.com/cm0109/Adoption_study.

For comparisons among species, ISR ASVs from the soft tissue swab/saliva samples were mapped to the updated Human Oral ISR database^5^. This database now constitutes of over 3,000 unique ISR sequences representing close to 300 different species of the most abundant oral bacteria and is publicly available for download (https://github.com/cm0109/ISR_database).

### Statistical analysis & visualizations

Statistical data analysis and visualizations were performed in RStudio version 1.1.456, with R version 3.5.1. ASV and species-OTU tables were rarefied using the R function *rrarefy* from the vegan^14^ package, using row minimums as subsample size. Relevant distance matrices (Bray-Curtis and Jaccard) were generated using the function *vegdist*, also from package vegan. Non-metric multidimensional scaling (NMDS) ordination was computed using the function metaMDS from the vegan package. Distribution of distances between two groups were compared using Wilcoxon rank sum test, with the function *stat_compare_means* from the package ggpubr^15^.

For statistical comparisons between groups where one of the groups consisted of paired subjects and the other of unrelated pairs, a permutation-based method was adopted such that correct distribution of test statistic could be obtained while accounting for dependency among pairs^16^. For this method, the observed test statistic was calculated as the median of the dissimilarity indices of a random subset of all possible unrelated pairs of subjects in the dataset minus the median dissimilarity of paired subjects. The size of the random subset was the same as the number of paired subjects so that the permutation test had comparable power as the Wilcoxon test used in other comparisons. The original pairing of subjects was then randomly permuted, and for each permutation, the new test statistic was calculated in the same way as for the original data. For each comparison, 1000 such permutations were computed, to obtain the empirical distribution of the test statistic. The two-sided p value was then calculated as the proportion of permutations where the absolute value of the test statistic is larger than or equal to the absolute value of the observed test statistic.

Multiple linear regression analysis was conducted using the function *lm* from package stats. Clinically relevant categorical variables recorded as none/mild/moderate/severe were converted to numerical representation for regression analysis using the scale 1/2/3/4 respectively. For measuring correlation, Spearman’s correlation test was applied, using function *cor.test* from package stats, which provided the p-value and Spearman’s rank correlation coefficient (rho) measure. Fisher’s exact test for metadata comparisons were performed using the R functions *fisher.test* from package stats.

R function *stars.pval* (package gtools v3.5.0) was used to convert numerical p-values to star notations. The convention used is as follows: if a p-value was less than 0.05, it was flagged with one star (*). If a p-value was less than 0.01, it was flagged with two stars (**). If a p-value was less than 0.001, it was flagged with three stars (***).

All visualizations were built using ggplot2 version 3.1.0^17^. Violin plots and box and whisker plots were generated using ggplot2 functions geom_violin and geom_boxplot, respectively. Venn diagrams were plotted using the *draw.pairwise.venn* function from the R package VennDiagram (ref). NMDS plots were constructed using *geom_points*, and vector fitting when applicable were drawn using *geom_segment*, both from ggplot2. Scatter plots were constructed using the ggplot2 function geom_points, smoothed using the LOESS fit smoothing by function geom_smooth (ggplot2). Bar plots were constructed using the ggplot2 function geom_bar (stat = “identity”).

## RESULTS

### Multi-niche, multi-resolution study framework

A multi-niche, multi-resolution approach **(Figure 1)** was implemented to compare oral microbial communities of children and adults, to determine mother to child bacterial transmission within genetically related and unrelated families. Subjects recruited for this study included 50 adoptive mother-child-pairs and 55 biological mother-child pairs. The adoptive group included only children adopted at birth by a non-genetic relative. The biological group was matched on age (p=0.29) and socioeconomic status to the adoptive group. Additionally, an extended family dataset of samples was obtained from 23 fathers and 16 siblings of the children in the biological group, allowing comparison of microbial profile similarity among siblings, couples and child-father pairs. Detailed meta-data including feeding and delivery mode, health measures, and demographics were also collected.

**Figure 1.**
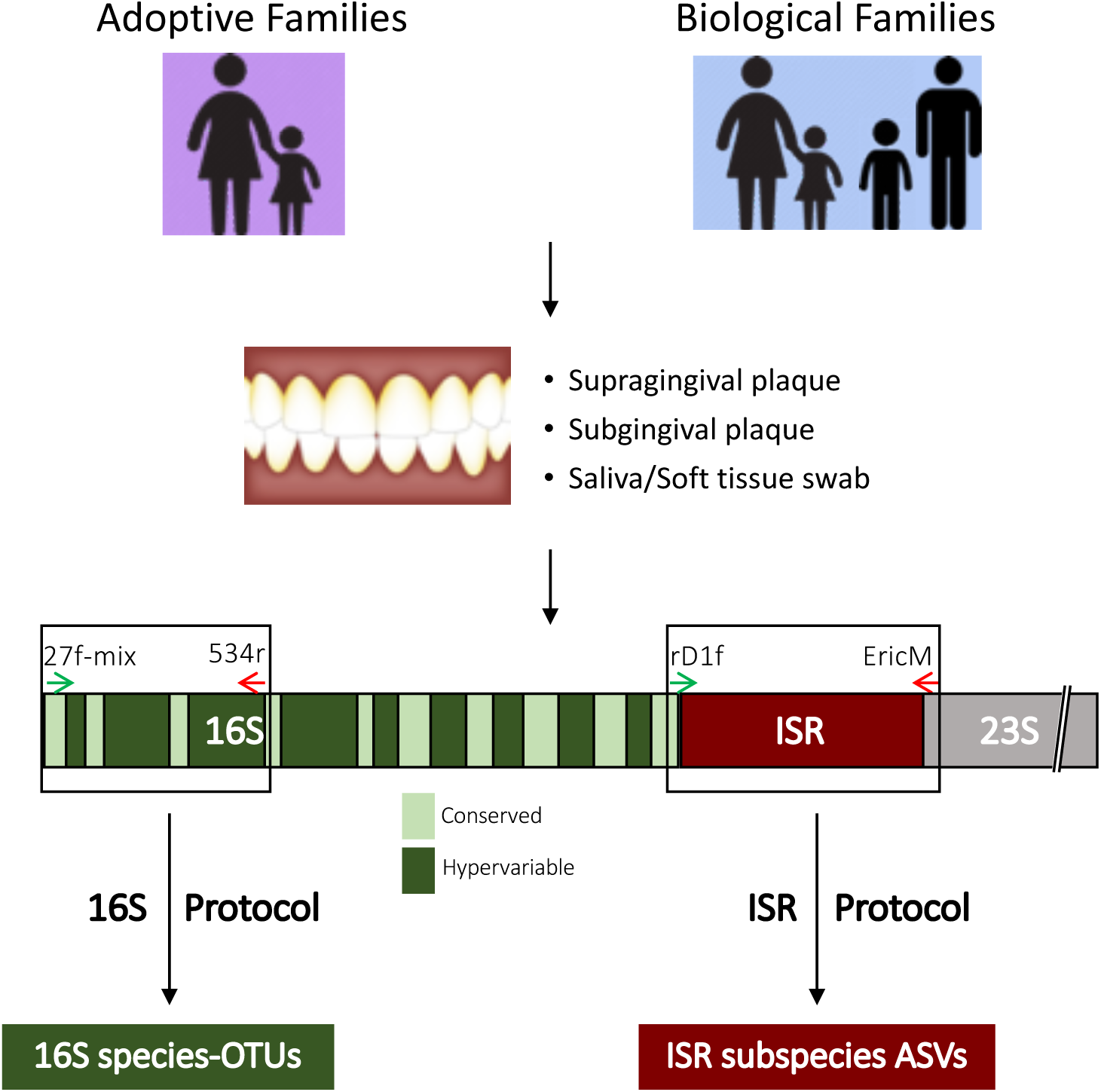
Overview of the multi-habitat and multi-resolution approach to compare microbial profiles. Adoptive and biological mother-child pairs were the main comparison groups. In addition, siblings and fathers in the biological group were recruited. Three distinct microbial habitats within the oral cavity were sampled – soft tissue and saliva, supragingival plaque, and subgingival plaque. For profiling species-level communities, amplicon sequencing targeting the 16S V1-V3 region was performed. For strain-level profiling, amplicon sequencing of the 16-23S Intergenic Spacer Region (ISR) was performed (see Methods).

To comprehensively profile microbial communities from multiple niches within the oral cavity, one soft tissue/saliva swab sample, one supragingival plaque sample, and one subgingival plaque sample were collected from each subject, with the exception of predentate children, from whom no tooth-associated sample could be collected. Microbial communities from each sample were independently profiled at both the species and strain level. Species level resolution was achieved through sequencing of the 16S V1-V3 region and mapping to the OSU CORE^12^ reference database of oral bacteria, and strain level resolution was achieved by targeted sequencing of the 16-23S Intergenic Spacer Region (ISR) combined with high-resolution processing using DADA2, as shown previously^5^.

For strain-level community analysis, the ISR-amplicon library was sequenced using the Illumina HiSeq 2500 platform and processed with DADA2^13^ to generate unique ISR amplicon sequence variants (ISR-ASVs) or ISR-type strains. Seven samples which did not generate sufficient sequences (cutoff 5000 reads) were excluded from analysis. A conservative approach was implemented in calling true biological strain variants by including in the analysis only those ASVs found in more than 1 sample. The final dataset consisted of 778 samples, representing 3,865 ASVs. This included 49 adoptive and 54 biological mother-child pairs for the saliva dataset, 46 adoptive and 53 biological pairs for the supragingival dataset, and 44 adoptive and 51 biological pairs for the subgingival dataset.

For the species-level community analysis, the same samples were used to generate a 16S V1-V3 amplicon library, sequenced on Illumina MiSeq platform, processed using Mothur^11^, and species level taxonomy was assigned to the reads using CORE^12^ oral 16S database. The final quality filtered dataset consisted of 709 samples, representing 581 species-level OTUs. This included 45 adoptive and 48 biological mother-child pairs for the saliva subset, 45 adoptive and 48 biological pairs for the supragingival subset, and 40 adoptive and 46 biological pairs for the subgingival subset.

Non-metric multidimensional scaling (NMDS) ordination using Bray-Curtis (BC) dissimilarities based on community membership was used to compare all the mother and child samples, both at the species (16S) and strain (ISR) levels (**Figure 2**). Profiling strain-level communities showed greater separation between the samples compared to species-level communities (**Supplementary Figure S1**).

**Figure 2.**
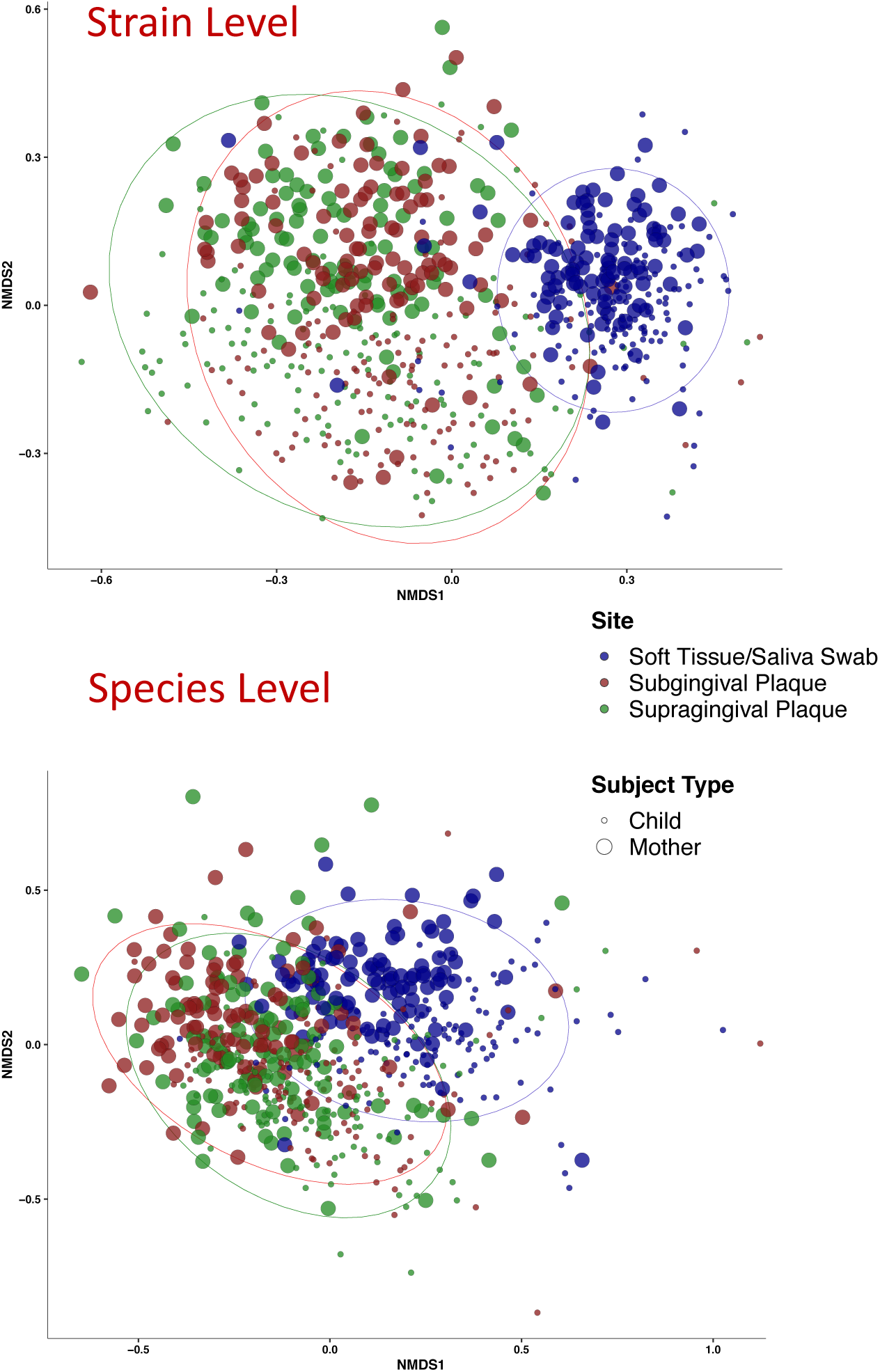
Beta-diversity comparison among samples by subject type and sampling site, at both species and strain levels. Non-metric multidimensional scaling (NMDS) plots using Bray-Curtis dissimilarities based on community membership, at ISR-strain level (top) and 16S Species level (bottom) are shown. At the lower resolution (species level) inter-sample distances are smaller as compared to strain level.

### No genetic influence on acquisition of oral bacteria

Similarity between the microbial profiles of adoptive and biological mother-child pairs was quantified using a distance-based approach. BC dissimilarities were computed for each mother-child pair based on presence/absence of species/strain variants. This index ranges from 0-1, with samples having exactly identical microbial communities scored at 0 and absolutely different communities scored at 1. We compared the BC distances between mother-child at both species and strain levels (**Figure 3**) using the Wilcoxon rank sum test. Microbial profiles of biological and adopted children were equally similar to their mothers, for all three niches we sampled: supragingival plaque, subgingival plaque and saliva/soft tissue. This was true at both species and strain level resolutions. BC dissimilarities of all possible unrelated mother-child pairings among the samples were also computed. A group containing all possible distances between any unrelated mother-child pairs is often compared with distances between adoptive or biological mother-child pairs using a Wilcoxon test or t-test, but this violates a basic assumption of the tests by re-using the same observations multiple times. Therefore, as an alternative to the widely used approach, we used a permutation-based method for statistical comparison between the unrelated group and the adoptive/biological groups (**see Methods for details**). At the level of strains, all mothers and their own children, regardless of genetic relationship, were significantly more similar to each other than unrelated mother-child pairs. This relationship was not as strong using the lower resolution species-level approach. Similar results were also observed using the Jaccard dissimilarity index (**ISR Swab/saliva dataset shown in Supplementary Figure S1**). Additionally, when using relative abundance measures in place of presence/absence, similar results for comparisons between adoptive/biological groups were observed (**ISR Swab/saliva dataset shown in Supplementary Figure S2**). Given that the 3 sites showed highly similar patterns, and the soft tissue swab/saliva provided the largest dataset for our comparisons because it included the predentate children, we used those samples as the primary dataset in subsequent analyses.

**Figure 3.**
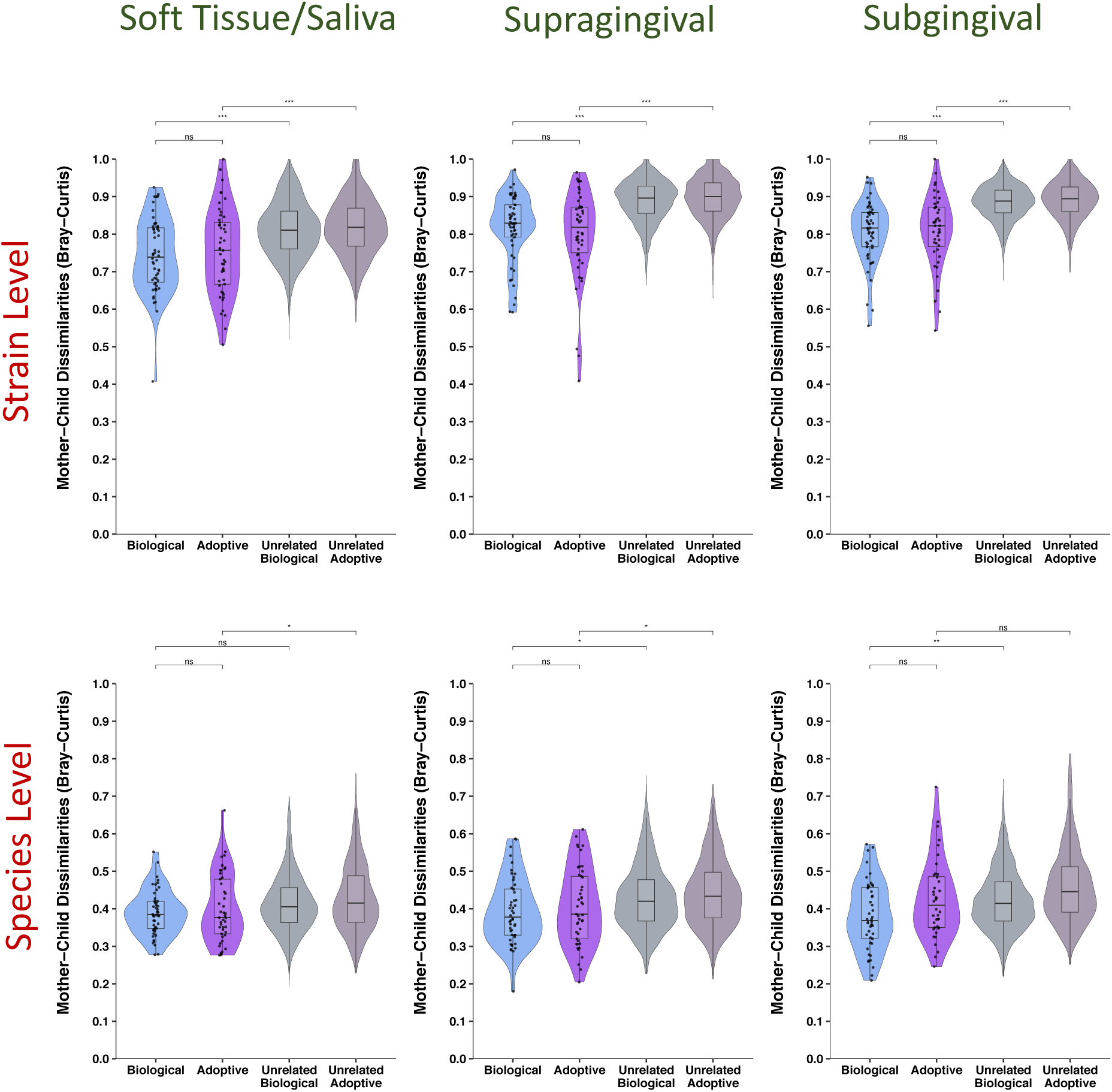
No influence of genetics on sharing of strains or species between mother and child. Violin plots with embedded box-and-whisker plots are shown here comparing the distribution of mother-child distances in the biological, adoptive, unrelated biological and unrelated adoptive groups, for the three sampling sites, at strain (**top panel**) and species level (**bottom panel**). No significant difference was observed in the mother-child dissimilarities for the biological and adoptive groups, at either species or strain levels, across the 3 distinct habitats within the oral cavity. Biological vs adoptive statistical comparisons were performed using Wilcoxon rank sum test, and related/unrelated comparisons were performed using a comparable permutation-based test (*See Methods*). Significance levels: ns: p > 0.05, * : p ≤ 0.05, **: p ≤ 0.01, ***: p ≤ 0.001.

We calculated the average number of species and strains shared between mother-child pairs for the soft tissue/saliva samples (**Figure 4**). Both adoptive and biological groups shared 44% of their microbiota at species level, and 15% at strain level. As expected, the fraction of shared species was much higher than fraction of shared strains, and unrelated mother-child pairs shared 4 times as many oral species as oral strains. Even though the set of species and strains shared between mothers and children made up 44% and 15% of the total *number* of species and strain variants, when considering *relative abundance*, they accounted for 93% and 48% of the total communities on average.

**Figure 4.**
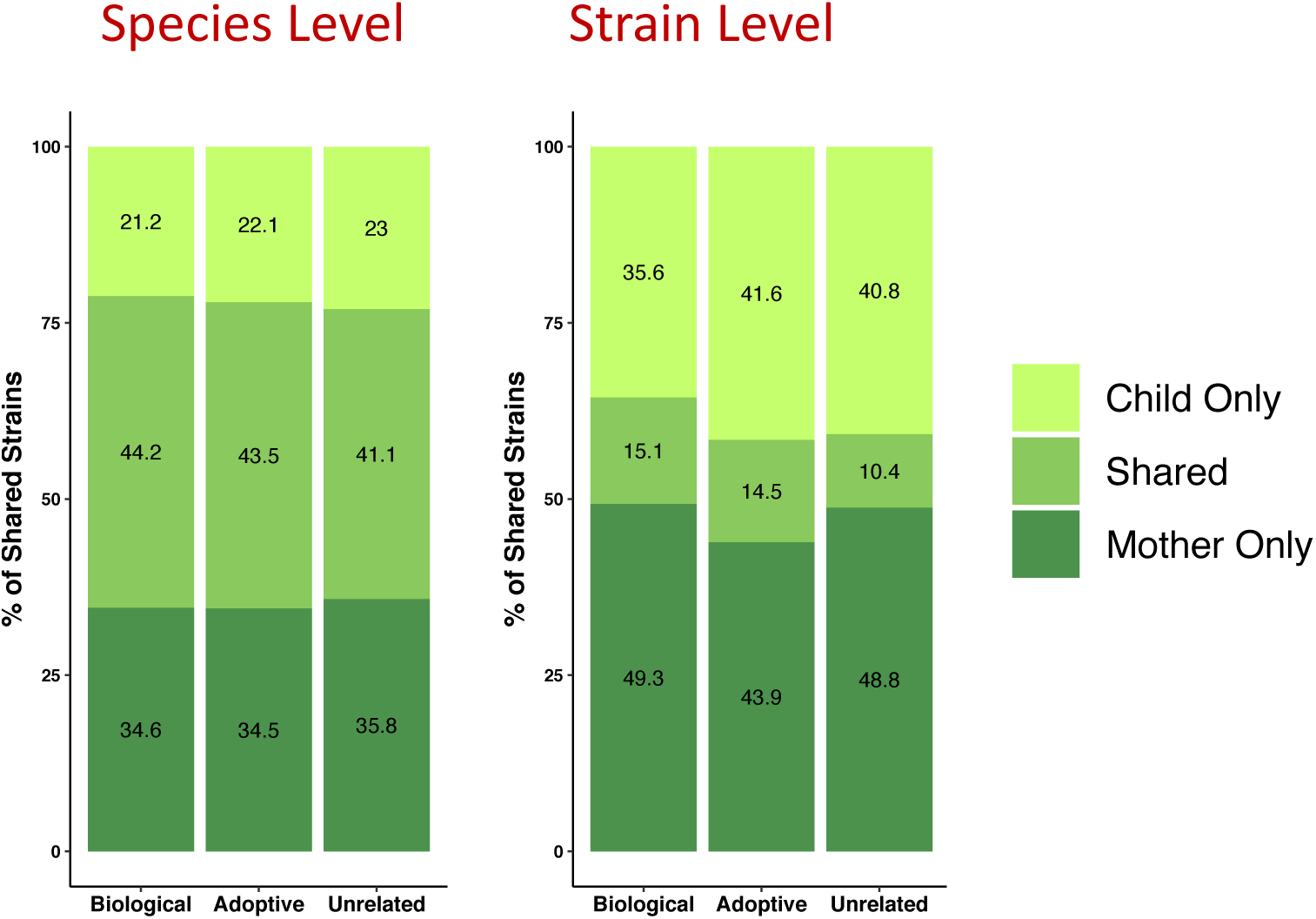
Adopted & Biological Mother-Child pairs shared similar percentage of species and strains. Stacked bar plots showing mean number of species OTUs/ISR strains shared by mother-child pairs in the adoptive, biological and combined unrelated groups for the saliva/soft tissue swab samples. The percentages of shared species/strains were comparable for the adoptive and biological groups at both resolutions

### Ruling out possible confounders including feeding and delivery mode

Extensive demographic and clinical data were collected for all subjects and examined to determine if any of these variables significantly influenced mother-child dissimilarities. Targeted recruitment lead to close matching of age, health factors, and socio-economic status between the biological and adopted children (no significant differences). However, the following 9 potentially confounding factors for the mother-child dissimilarities were found to be significantly different between the two groups by Wilcoxon rank sum or Fisher’s exact test: child’s feeding mode, child’s gingivitis level, child’s race, child’s tongue biofilm level, mother-child race match, mother’s age, mother’s plaque level, mother’s tongue biofilm level and mother’s gingivitis level. To assess the confounding effect, a univariate regression analysis with mother-child distance as the dependent variable and group as the independent variable was performed, and a multiple regression analysis with the unbalanced factors as additional independent variables was also performed. The estimated regression coefficient of the group variable changed substantially from the univariate model (beta= -0.017, p=0.433) to the multiple regression model (beta=-0.056, p=0.226). However, the group variable remained nonsignificant even after adjusting for the confounding effect in the multiple regression model, thereby ruling out any significant confounding effect on the mother-child dissimilarities. Within the biological group neither feeding mode nor delivery mode had a significant impact on mother-child distances for any niche (**Supplementary Figure S3**).

### Extended family comparisons also fail to show influence of genetics

An extended dataset of the biologically related families that included fathers and siblings was analyzed to further explore the relative contributions of genetics and shared environment. Comparisons of distance between various related and unrelated pairs of subjects for the soft tissue/saliva dataset at strain level is shown in **Figure 5**. Cohabitating mother-father couples and sibling pairs showed the greatest oral microbiota resemblance of any pairing examined, sharing significantly more strains than mother-child or father-child pairs. Mothers and fathers were equally similar to their children, and both mother- and father-child pairs were significantly more similar than unrelated mother- and father-child pairings. Technical replicates consisting of samples that were processed through the ISR pipeline in duplicates were highly similar. Analysis of the extended family samples from the subgingival and supragingival dataset also produced similar results (**Supplementary Fig S4**). Taken together, these findings provide further evidence that shared environment and contact, rather than genetic background, is the primary determinant of microbial community structure.

**Figure 5.**
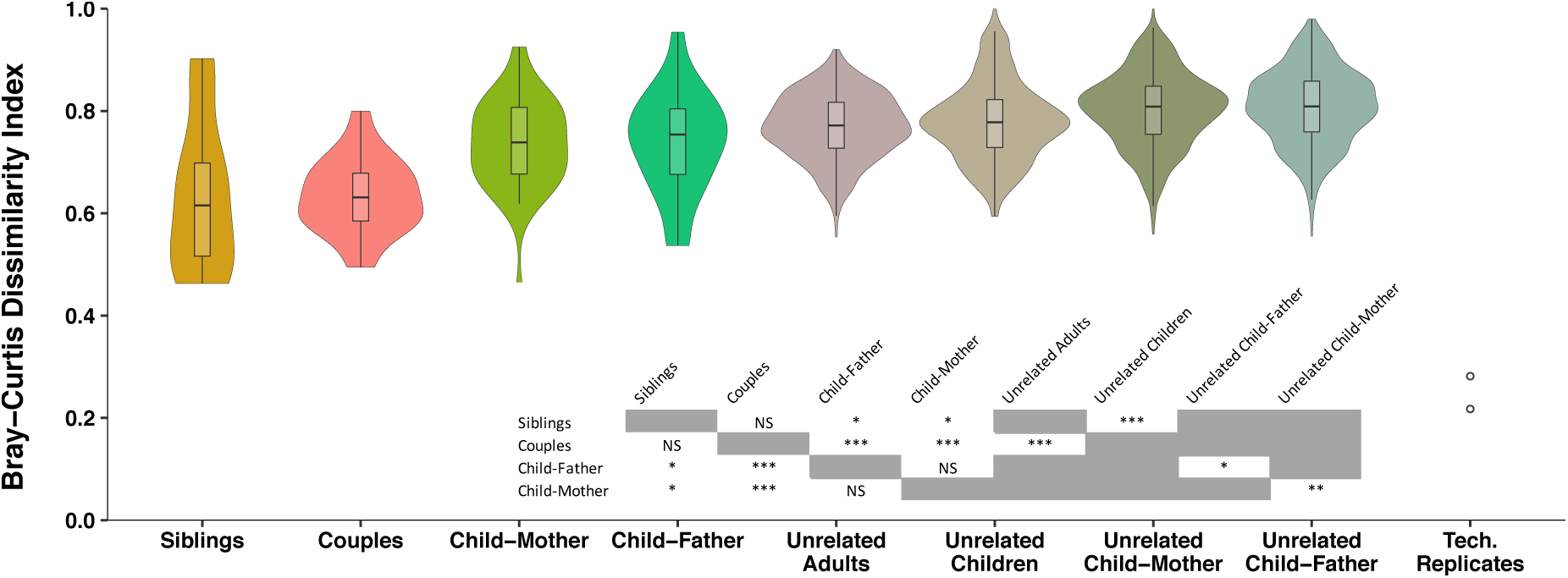
Shared environment led to greater similarity between individual’s oral microbial communities at strain level. Comparing microbial community similarities among different family groups, based on saliva/soft tissue swab samples from the extended biological family dataset. Dissimilarities between individuals were lowest among cohabitating couples and siblings, even compared to child-mother and child-father. Children’s oral microbiota were equally similar to their fathers as they were to their mothers. Couples and siblings were more similar to themselves, compared to unrelated adults and unrelated children, respectively. Technical replicates were highly similar to one another. Statistical comparisons were performed using Wilcoxon rank sum test and the previously used permutation test (when including unrelated groups).

### Older children’s microbiota more similar to their mothers

Child’s age is known to be a major determinant of oral microbiota composition, but targeted recruitment allowed us to ensure that the child’s ages in the adoptive and biologic group were balanced (**Fig 6a**). Since our results showed that the biological and adoptive children were equally similar to their mothers, the two groups were combined and examined to determine if child’s age had an effect on mother-child dissimilarities. A total of 101 children were considered for this analysis, with an age range of 3 months to 6 years. Contrary to our hypothesis, we saw that younger children’s microbial communities were less similar to that of their mothers compared to older children (**Fig 6b**). The strong negative association between child’s age and mother-child dissimilarities (Spearman correlation coefficient rho = -0.33, p=0.002) lead us to further explore this relationship. We hypothesized that increasing overall diversity of the oral microbial communities with age of the child may be responsible for increasing similarity of children’s microbiota to their mothers with age. Indeed, our results show that the Shannon Diversity Index, a measure of alpha diversity, increased sharply during the early years, reaching levels comparable to those of the mothers for most children by age 5 years (**Fig 6c**).

**Figure 6.**
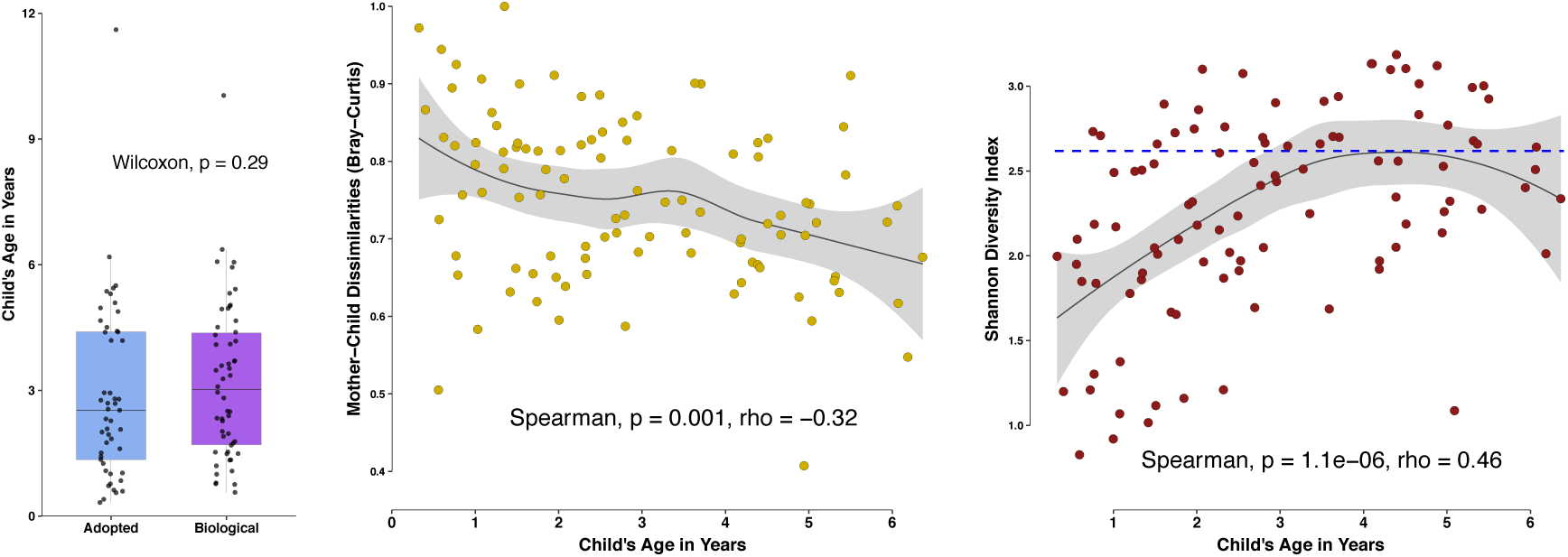
Child’s age was a significant determinant of mother-child dissimilarities. **a.** Box and whisker plot showing distribution of ages among the biologic and adoptive group children. **b.** Scatterplot exhibiting the relationship of mother-child distances with age of the child. A strong negative correlation between mother-child dissimilarities and child’s age was observed. Both adoptive and biologic children were included in this analysis, and two older children (>=10 years) were excluded (n=101). **c.** For the same 101 children, the alpha diversity measure Shannon Diversity Index was plotted against age. The blue dotted line represents mean Shannon Diversity for mothers of those children. Alpha diversity also showed strong positive correlation with child’s age, and most older children’s diversities were similar to the adults. Strength and direction of associations were measured using Spearman’s rank-order test. Scatter plots were smoothed using the regression method LOESS fit. Analysis was based on strain level communities.

### No influence of genetics seen for individual species

While considering the entire bacterial community as a whole did not show any effect of genetic kinship, we wanted to explore whether individual species of bacteria showed differences in degree of strain matching between biologic and adoptive mother-child pairs. Our extensive database of oral ISR sequences allowed us to assign species level taxonomy to ISR-type strains, and a list of the most abundant species, along with the number of strains identified for each, is shown in **Supplementary Table ST1**. To determine if fidelity of transmission varied for individual oral bacteria species, we compared the adoptive and biologic mother-child dissimilarities from the saliva/soft tissue swab dataset for the ten most abundant species in the saliva/soft tissue swab dataset (**Figure 7**). No species showed any significant difference between the adoptive and biologic groups, suggesting no differential heritability.

**Figure 7.**
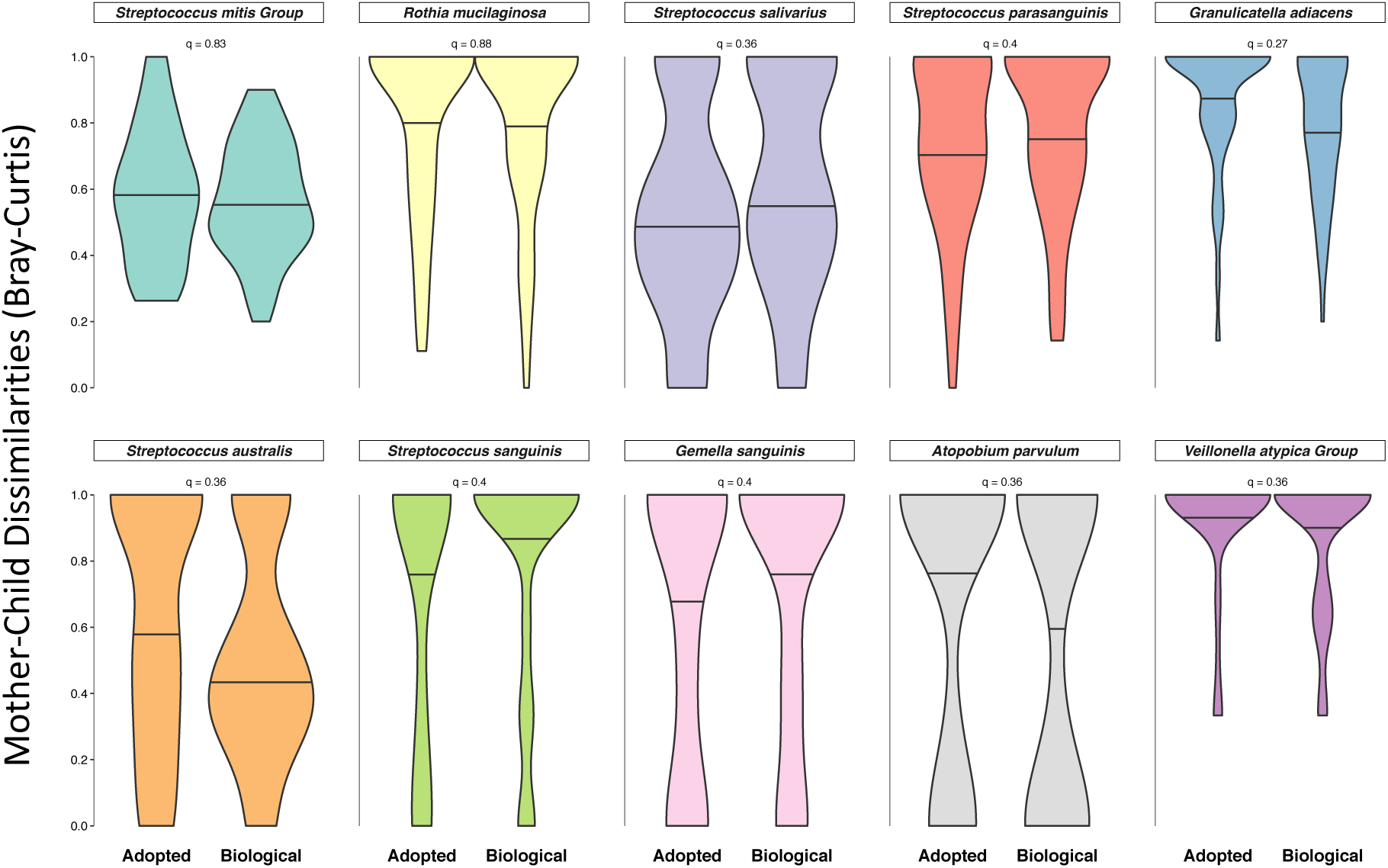
No influence of genetics on mother-child distances for individual species. Violin plots comparing the distribution of mother-child dissimilarities in the biologic and adoptive groups. For the 10 most abundant species, Bray-Curtis dissimilarities were generated based on presence/absence of strains for each species. Mother-child distances (dissimilarities) were not significantly different between the adoptive and biologic groups for any species. Wilcoxon rank sum test was used for statistical comparisons, and p-values generated were corrected for false positives (Benjamini-Hochberg procedure) to generate q-values shown. The 50^th^ quantile of each distribution is marked for comparison. Data is based on the saliva/soft tissue swab samples.

## DISCUSSION

Most investigations of the relative contribution of genetics and environment to the human microbiome have used a monozygotic vs dizygotic twin model. To our knowledge, this is the first study to directly investigate parent-to-child microbial transmission by using an adoptive *vs* biologic mother-child study design. Only children adopted by a genetically unrelated family were included. The genetic distances between African and northern European populations are among the greatest found in modern humans^18^, and many of our adoptive dyads were composed of white mothers and African-American children, providing maximum genetic distance between unrelated. Only children adopted at birth were included to minimize contact with the birth mother’s microbiota. Thus, our adoptive vs biologic study design allowed us to directly explore the contribution of host genetics to acquisition of oral microbiota. We found no measurable effect of genetic relatedness on how closely children’s oral microbiota resembled that of their mothers (**Figure 2**). These results suggest an alternative interpretation for the findings of two twin-based studies that have shown a small but significantly greater microbiota similarity between MZ than DZ twins^19,20^. What MZ twins do share to a greater extent than DZ twins and adopted siblings are important environmental determinants such as greater social closeness and intimacy, more similar treatment from others, and greater tacit coordination in choice making^21^. We suggest that the greater similarity of oral microbial communities previously observed in MZ twins as compared to DZ twins may be mediated through shared environmental factors and not a direct effect of genetically determined host factors.

Although little previous evidence is available at the strain level, studies at species level have provided evidence that shared environment is important in shaping oral microbiota composition. In one of three published oral microbiome twin studies^19,20,22^, MZ twin pairs were found to be no more similar than DZ twin pairs, and they became less similar when they no longer cohabited^22^. Kissing couples were observed to share highly similar tongue dorsum microbial communities^23^, and shared household was more important than genetic relationship in a dataset from an extended family^24^. Another study found that household members, particularly couples, shared more of their microbiota than individuals from different households^25^. While these studies did not resolve microbial communities at strain level, and thus did not provide sufficient resolution to track transmission, they support our finding that shared environment and direct contact are the drivers of microbial community structure.

Findings from pairings within our extended biological family dataset that included fathers and siblings further supported the importance of environment over genetics in determining oral microbial community composition (**Figure 5**). The pairings that showed the greatest similarity were sibling pairs and married couples, and they were equally similar despite the difference in genetic relatedness. Parent-child pairings were less similar than spouses or siblings, again pointing to degree of contact and age-related factors as environmental determinants of microbial similarity.

Common biologic features of mother-child relationship have been previously investigated and have been found to have some effect on microbial communities at the species level^26,27^. We saw no effect on the fidelity of mother-child strain sharing from vaginal birth or breast feeding (**Supplementary Fig S4**). We also observed that mother-child similarity was greater in older children **(Figure 6).** In addition, father-child oral microbiota matched just as closely as that of mothers and children (**Figure 5**). Together these observations suggest that any impact of breast feeding or delivery mode on oral strain sharing is negligible relative to other environmental factors.

Subgingival and supragingival niches have distinct ecologies and microbial community profiles, and dysbiosis of these communities causes the two major oral diseases, dental caries and periodontitis^28^. The saliva and soft-tissue surfaces, being easily accessible, have been most commonly sampled, but do not reflect the disease-associated communities of greatest interest. Due to the importance of biogeographic diversity within the oral cavity^29^, we sampled three distinct sites–soft tissue/saliva surface, supragingival plaque biofilm and subgingival plaque biofilm, and confirmed that the lack of a measurable effect of genetics on microbial communities was consistent across all three niches.

Species-level microbial identification using 16S rRNA gene sequencing provides limited power to track bacterial transmission, although it may provide a good indicator of functional similarity of bacterial communities. Our previous work showed that individuals have relatively similar microbial profiles when analyzed at the species level, but their microbiota are distinct and personalized at the subspecies level^5^. For this study, we used both 16S species-level and ISR strain-level sequencing approaches to compare microbial profiles. Neither approach detected a difference between adoptive and biologic dyads, but our findings illustrate the greater power of the ISR-based approach to distinguish microbial communities (**Figure 3**). For example, the level of strain sharing was quite low between random pairings, only 10 percent, while at the species level it was 41 percent (**Figure 4**). Only the strain-level analysis consistently able to detect differences between mother-child and random pairings.

Previous studies using relative abundance data and interclass correlation coefficient (ICC) and ACE modeling in twins have suggested that some taxa might be more “heritable” than others, but the findings have not been consistent across studies^19,20^. Strain-level resolution achieved in our study allowed us to more directly address variation in heritability among species. We compared the frequency of strain concordance between biological and adoptive parents and offspring for the most common species (figure 7). Depth of sequencing limited analysis to the 10 most abundant species. None of these showed significantly different mother-child distances when comparing adoptive to biologic groups, indicating no effect of host genetics for any species (**Figure 7**).

The host is clearly active in shaping the composition of the oral microbiome, since fewer than a thousand of the many bacterial species in the larger environment are capable of colonizing the human oral cavity. Our findings suggest that these control mechanisms are universally shared among humans, since no effect of genetic relatedness on fidelity of microbial transmission could be detected. Instead our findings point towards contact and shared environment being the driving factors of microbial transmission, with a unique combination of these factors ultimately shaping a highly personalized human oral microbiome.

## Acknowledgements

Funding for this research study was provided by the National Institute of Dental and Craniofacial Research of the U.S. National Institutes of Health, under the award number R01-DE024327. We are extremely grateful to the local adoption agencies that helped us connect with adoptive families willing to participate in this research. We are also grateful to Center of Science and Industry (COSI) Columbus for assistance with recruitment of the biologically related families.

We are grateful for assistance in the laboratory from Haella Holmes, Laura Mason and Laura Cook, assistance in bioinformatic processing from Thomas Ellison and Zach Thomson, and overall guidance on this project from Clifford Beall.

## Author Contributions

ALG and EJL conceived the project idea and ALG designed the study. COM lead the recruitment of the adoptive families and HMS lead the recruitment and sampling of the biological families, with assistance from SBH, under the guidance of ALG. SBH and CM were responsible for the sequencing library preparation, with guidance from EJL. CM performed the bioinformatic processing and statistical data analysis, with inputs from XG, and guidance from AN and ALG. CM and ALG prepared the manuscript.

## Notes

https://github.com/cm0109/Adoption_study

## BIBLIOGRAPHY

1. Huttenhower, C. et al. Structure, function and diversity of the healthy human microbiome. Nature 486, 207–214 (2012).

2. Dewhirst, F. E. et al. The human oral microbiome. J. Bacteriol. 192, 5002–5017 (2010).

3. Sulyanto, R. M., Thompson, Z. A., Beall, C. J., Leys, E. J. & Griffen, A. L. The Predominant Oral Microbiota Is Acquired Early in an Organized Pattern. Sci. Rep. 9, (2019).

4. Lloyd-Price, J. et al. Strains, functions and dynamics in the expanded Human Microbiome Project. Nature 550, 61–66 (2017).

5. Mukherjee, C., Beall, C. J., Griffen, A. L. & Leys, E. J. High-resolution ISR amplicon sequencing reveals personalized oral microbiome. Microbiome 6, 153 (2018).

6. Faith, J. J. et al. The long-term stability of the human gut microbiota. Science (80-.). 341, 1237439–1237439 (2013).

7. Ramos-Gomez, F. & Ng, M.-W. Into the future: keeping healthy teeth caries free: pediatric CAMBRA protocols. J. Calif. Dent. Assoc. 39, 723–33 (2011).

8. Si, J., Lee, S., Park, J. M., Sung, J. & Ko, G. P. Genetic associations and shared environmental effects on the skin microbiome of Korean twins. BMC Genomics 16, 992 (2015).

9. Zaura, E., Keijser, B. J., Huse, S. M. & Crielaard, W. Defining the healthy ‘core microbiome’ of oral microbial communities. BMC Microbiol. 9, 259 (2009).

10. Weisburg, W. G., Barns, S. M., Pelletier, D. A. & Lane, D. J. 16S ribosomal DNA amplification for phylogenetic study. J. Bacteriol. 173, 697–703 (1991).

11. Schloss, P. D. et al. Introducing mothur: Open-source, platform-independent, community-supported software for describing and comparing microbial communities. Appl. Environ. Microbiol. 75, 7537–7541 (2009).

12. Griffen, A. L. et al. CORE: A phylogenetically-curated 16S rDNA database of the core oral microbiome. PLoS One 6, 1–10 (2011).

13. Callahan, B. J. et al. DADA2: High-resolution sample inference from Illumina amplicon data. Nat. Methods 13, 581–583 (2016).

14. Oksanen, J. et al. vegan: Community Ecology Package. (2018).

15. Kassambara, A. ggpubr: ‘ggplot2’ Based Publication Ready Plots. (2018).

16. Reiss, P. T., Stevens, M. H. H., Shehzad, Z., Petkova, E. & Milham, M. P. On distance-based permutation tests for between-group comparisons. Biometrics 66, 636–643 (2010).

17. Wickham, H. et al. ggplot2: Create Elegant Data Visualisations Using the Grammar of Graphics. (2019).

18. Tierney, B. T. et al. The Landscape of Genetic Content in the Gut and Oral Human Microbiome. Cell Host Microbe 26, 283-295.e8 (2019).

19. Gomez, A. et al. Host Genetic Control of the Oral Microbiome in Health and Disease. Cell Host Microbe 22, 269-278.e3 (2017).

20. Demmitt, B. A. et al. Genetic influences on the human oral microbiome. BMC Genomics 18, 1–15 (2017).

21. Segal, N. L., McGuire, S. A., Miller, S. A. & Havlena, J. Tacit coordination in monozygotic twins, dizygotic twins and virtual twins: Effects and implications of genetic relatedness. Pers. Individ. Dif. 45, 607–612 (2008).

22. Stahringer, S. S. et al. Nurture trumps nature in a longitudinal survey of salivary bacterial communities in twins from early adolescence to early adulthood. Genome Res. 22, 2146–2152 (2012).

23. Kort, R. et al. Shaping the oral microbiota through intimate kissing. Microbiome 2, (2014).

24. Shaw, L. et al. The human salivary microbiome is shaped by shared environment rather than genetics: Evidence from a large family of closely related individuals. MBio 8, e01237–17 (2017).

25. Song, S. J. et al. Cohabiting family members share microbiota with one another and with their dogs. Elife 2013, 1–22 (2013).

26. Holgerson, P. L. et al. Oral microbial profile discriminates breast-fed from formula-fed infants. J. Pediatr. Gastroenterol. Nutr. 56, 127–136 (2013).

27. Lif Holgerson, P., Harnevik, L., Hernell, O., Tanner, A. C. R. & Johansson, I. Mode of birth delivery affects oral microbiota in infants. J. Dent. Res. 90, 1183–1188 (2011).

28. Wade, W. G. The oral microbiome in health and disease. 69, 97–114 (2015).

29. Mark Welch, J. L., Dewhirst, F. E. & Borisy, G. G. Biogeography of the Oral Microbiome: The Site-Specialist Hypothesis. Annu. Rev. Microbiol. 73, 335–358 (2019).

