## Supplemental Material for "Acquisition of Oral Microbiota is Driven by Environment, Not Host Genetics"

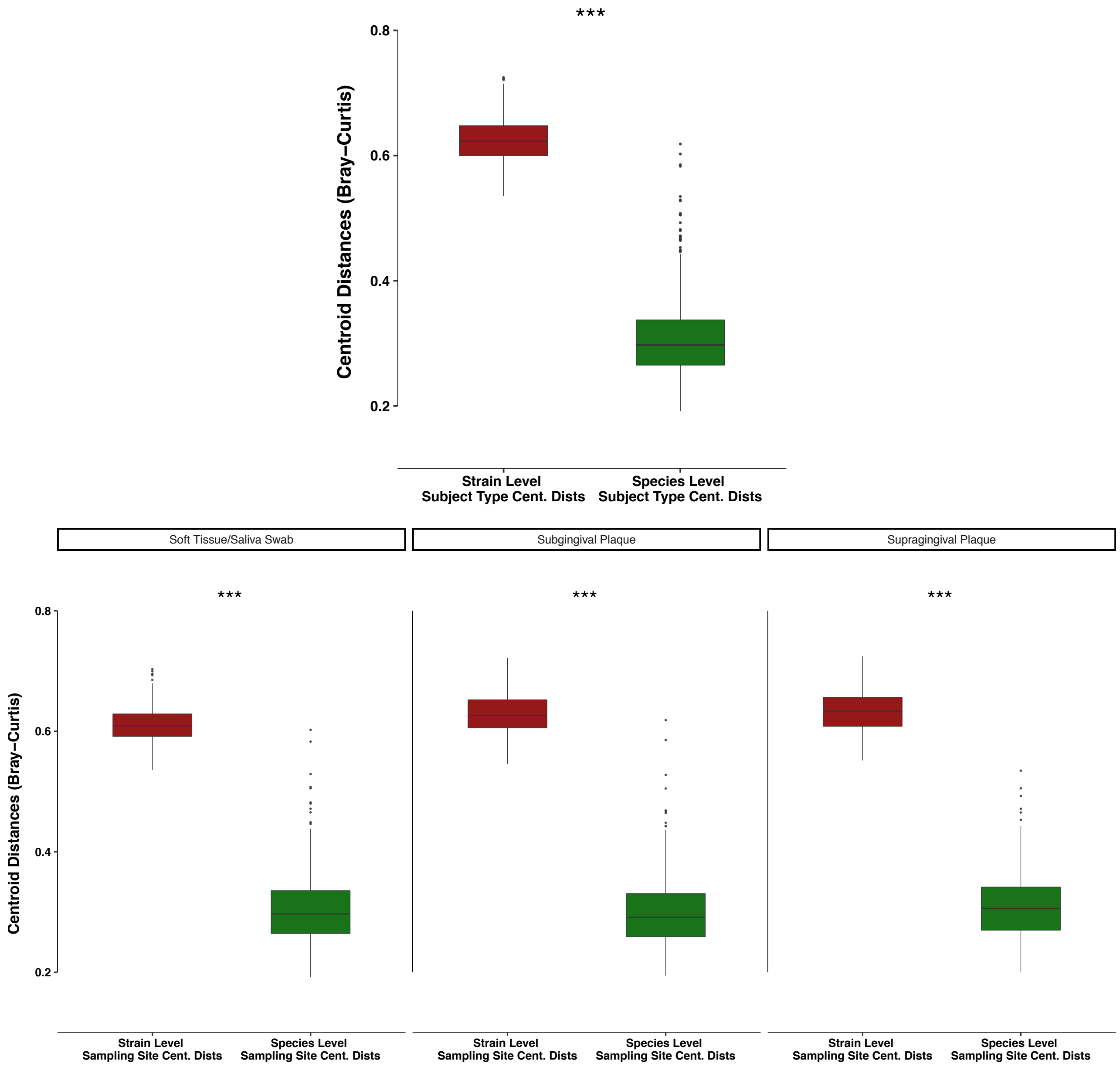


**Figure S1. Strain-level community characterization led to increased separation between samples.** Comparison of centroid distances for strain and species level communities, both in terms of subject type (mother/child) separation and sampling site (saliva/swab, subgingival and supragingival plaque) separation showed that samples were significantly better separated at the strain level. P-values were generated using paired Wilcoxon rank sum test (significance level *** refers to $p < 0.001$).

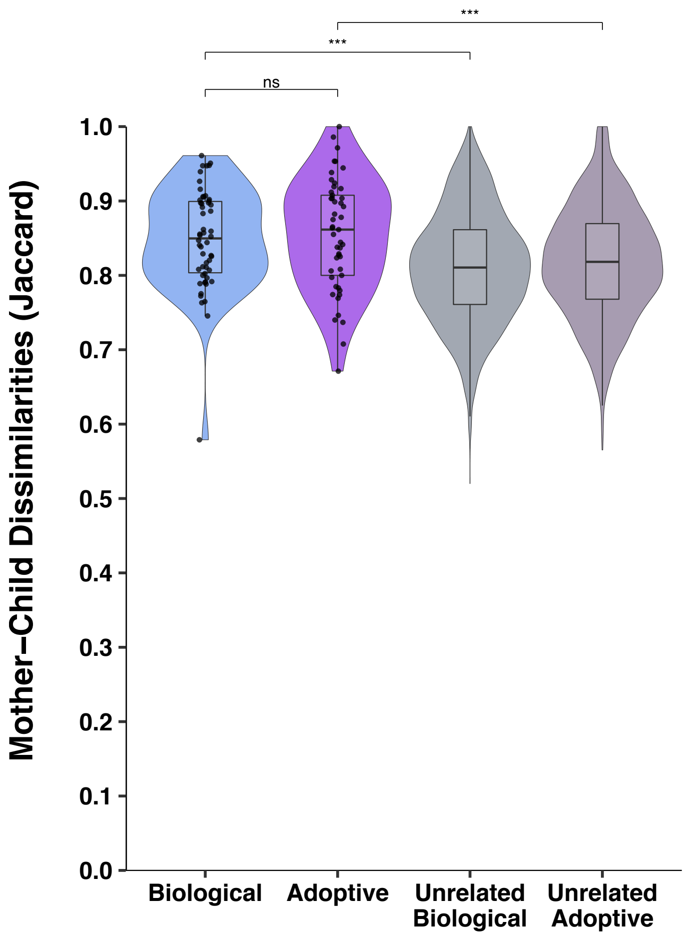


**Figure S2. No influence of genetics on sharing of strains between mother and child using Jaccard dissimilarities.** The saliva/soft tissue swab samples were also analyzed using the **Jaccard dissimilarity** indices computed based on presence/absence of ISR strains. The results were very similar to what was obtained using Bray-Curtis dissimilarities. No significant difference was observed in the mother-child dissimilarities between the biologic and adoptive groups, and both biologic and adoptive children's oral microbiota were significantly more similar to their own mothers' than unrelated mothers Distribution of distances are shown using violin plots, with embedded box and whisker plots. Biological vs adoptive statistical comparisons were performed using Wilcoxon rank sum test, and related/unrelated comparisons were performed using the previously described permutation test.

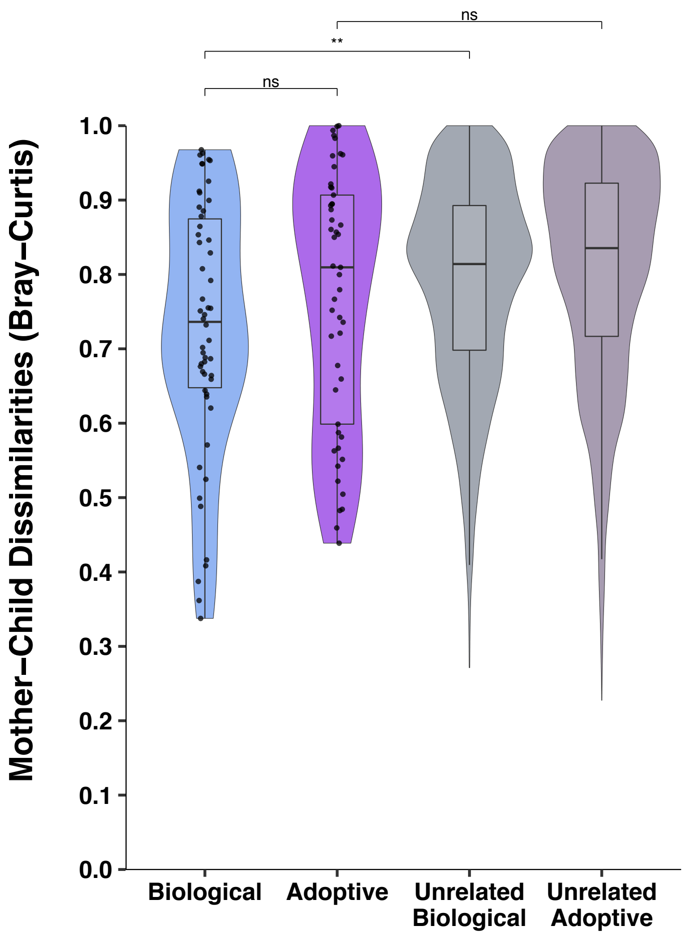


**Figure S3. No influence of genetics on sharing of strains between mother and child using relative abundance of stains.** Bray-Curtis dissimilarities between the mother-child pairs for the Saliva/soft tissue swab samples were also computed based on **relative abundance of ISR strains**. No significant difference was observed in the mother-child dissimilarities between the biologic and adoptive groups. While the biological group children's oral microbiota was significantly more similar to their own mothers' than unrelated mothers, the same distinction could not be made for the adoptive group. Biological vs adoptive statistical comparisons were performed using Wilcoxon rank sum test, and related/unrelated comparisons were performed using the previously described permutation test.

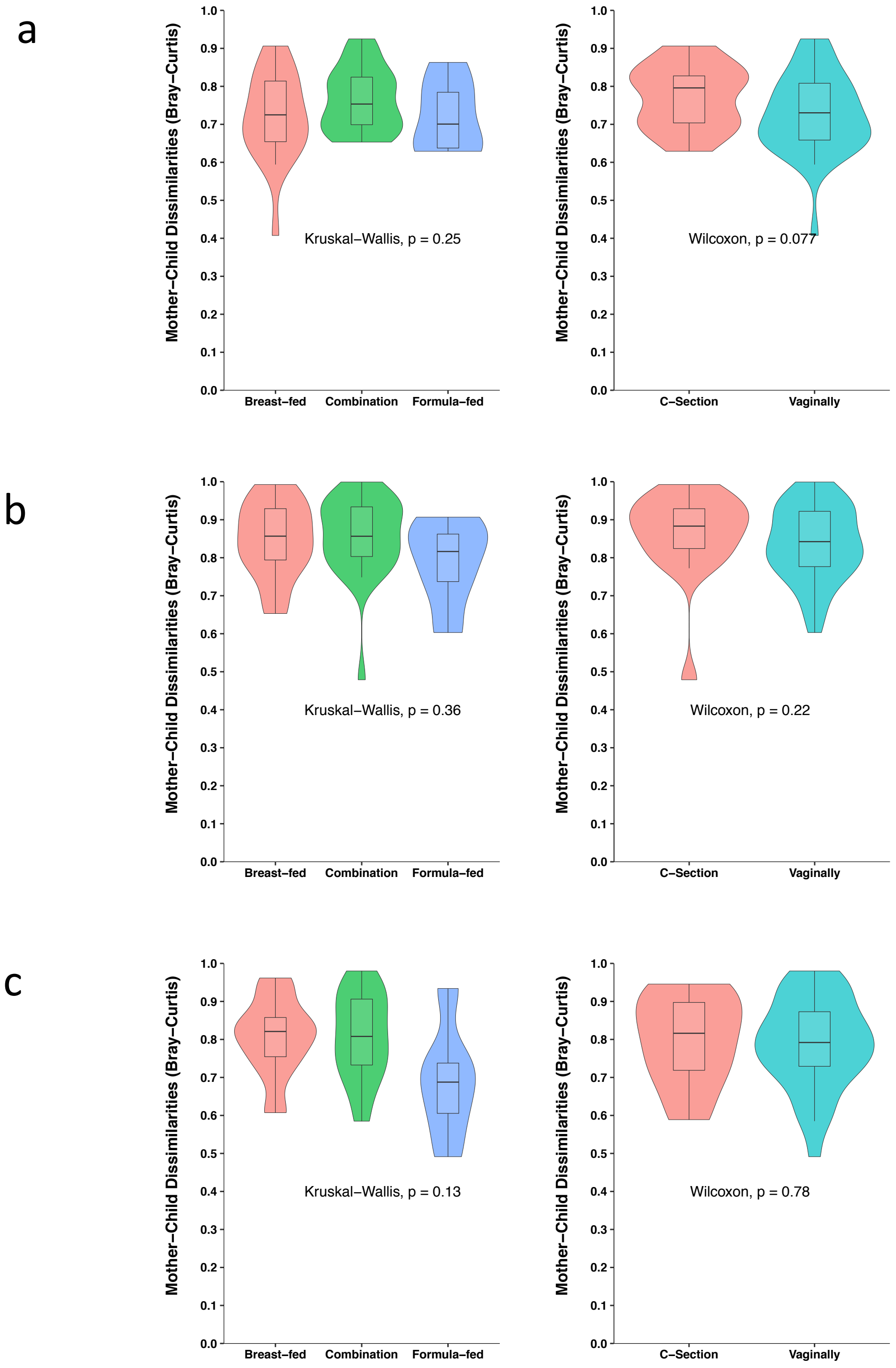


**Figure S4. Effect of feeding and delivery modes on mother-child distances.** Differences in feeding mode (right) or delivery mode (left) among the biological group children did not have any significant effect on the mother-child dissimilarities, for either the **a)** saliva/soft tissue swab, **b)** supragingival or **c)** subgingival plaque samples.

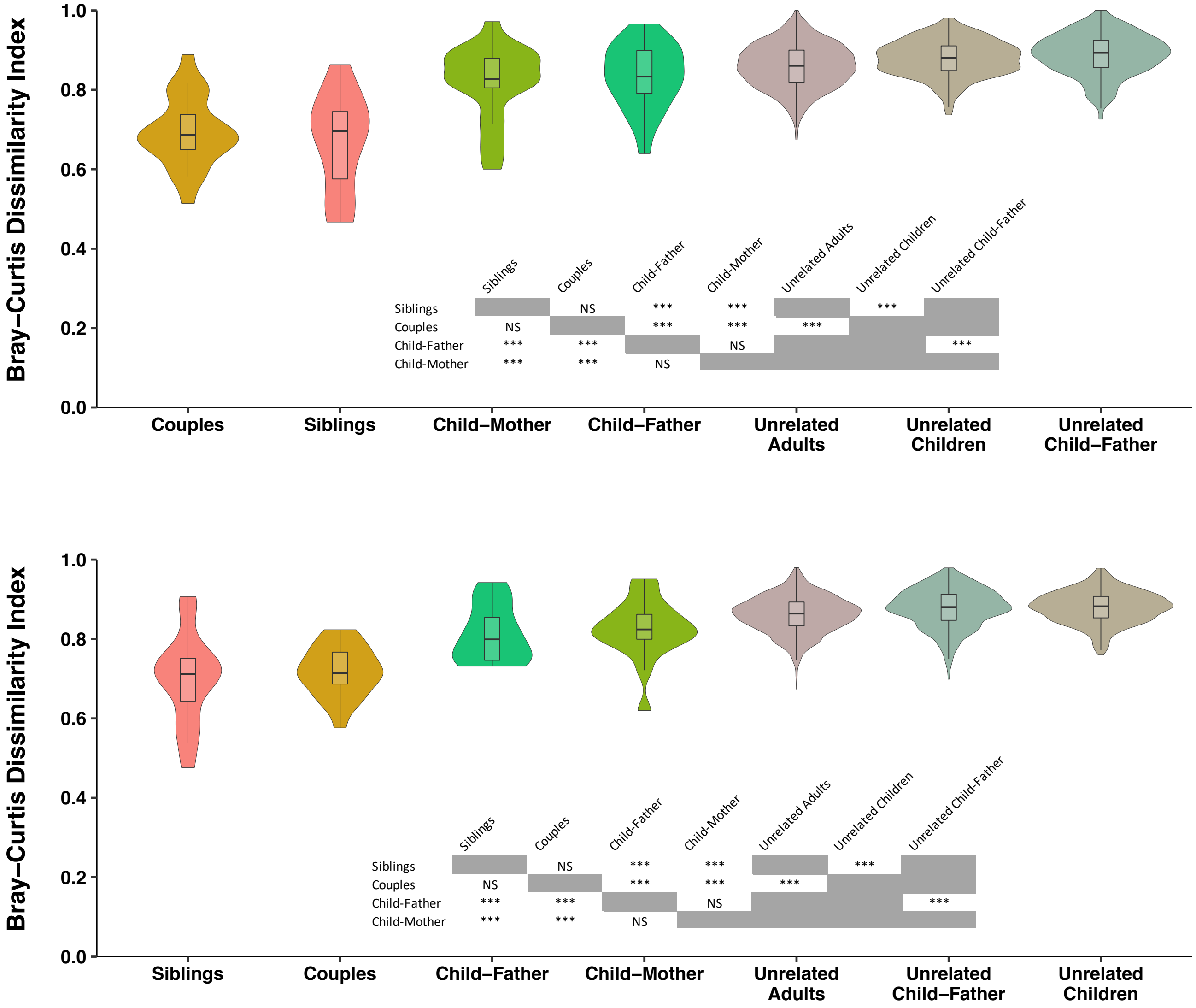


**Figure S5. Extended family comparisons using plaque samples show results similar to saliva samples.** Comparing microbial community similarities among different family groups, based on **supragingival plaque** (**top**) and **subgingival plaque** (**bottom**) samples from the extended biological family dataset. Shared environment/contact lead to greater oral microbiota composition similarity, and no evidence of genetic influence was detected. Statistical comparisons were performed using Wilcoxon rank sum test and a custom permutation test (when including unrelated groups).

| Species OTU | # of ISR-Type Strains | Total Seq. Counts | % of Subjects |
|---|---|---|---|
| *Streptococcus mitis pneumoniae infantis oralis* | 183 | 3648025 | 100 |
| *Rothia mucilaginosa* | 203 | 1323837 | 98.95 |
| *Streptococcus vestibularis salivarius* | 32 | 364929 | 95.44 |
| *Streptococcus parasanguinis* | 85 | 364200 | 95.44 |
| *Granulicatella adiacens* | 200 | 307273 | 98.95 |
| *Streptococcus australis* | 11 | 166784 | 83.16 |
| *Streptococcus sanguinis* | 44 | 115959 | 82.46 |
| *Gemella sanguinis* | 30 | 95432 | 88.77 |
| *Atopobium parvulum* | 9 | 52451 | 72.98 |
| *Veillonella atypica dispar parvula* | 103 | 43473 | 83.51 |
| *Neisseria meningitidis polysaccharea* | 13 | 42371 | 59.3 |
| *Moraxella osloensis* | 116 | 38833 | 46.67 |
| *Streptococcus gordonii* | 16 | 39042 | 35.79 |
| *Haemophilus parainfluenzae* | 43 | 31421 | 68.77 |
| *Gemella haemolysans* | 46 | 26062 | 48.07 |
| *Streptococcus cristatus* | 14 | 16893 | 43.86 |
| *Gemella morbillorum* | 21 | 15446 | 30.53 |
| *Rothia aeria* | 28 | 14582 | 46.32 |
| *Rothia dentocariosa* | 28 | 9174 | 19.65 |
| *Streptococcus intermedius constellatus* | 10 | 7077 | 16.84 |

**Table ST1. List of the 20 most abundant oral bacteria species.** Data is based on saliva/soft tissue swab samples.
